# Framed RSA: Representational comparisons that honor both geometry and population-mean response preferences

**DOI:** 10.1101/2025.07.10.664257

**Authors:** JohnMark Taylor, Heiko Schütt, Nikolaus Kriegeskorte

## Abstract

Representational similarity analysis (RSA) characterizes the geometry of neural activity patterns elicited by different stimuli while discarding information about neural response preferences, regional population-mean activity, and the absolute location and orientation of the patterns in the multivariate response space. When evaluating alternative representational models, invariance to certain aspects of the neural code is desirable because systems might use superficially different encodings to implement the same computations. However, neural preferences and regional-mean activation are arguably physiologically and mechanistically important, and so we may want our models to predict them correctly. Here we introduce a novel analysis technique, framed RSA, which honors both geometry and population-mean preferences in evaluating model-predicted representations. To achieve this, we augment the set of patterns that define the geometry by two reference patterns: the all-zero point (origin) and an all-*c* (uniform constant) pattern in the multivariate response space, enabling RSA to incorporate information about the profile across stimuli of regional population-mean activations and about the global location and orientation of the ensemble of response patterns. We show that framed RSA improves model-selection accuracy when ground truth is known, considering brain-region identification (using human fMRI data from the Natural Scenes Dataset and macaque intracranial recording data from the Things Ventral Stream Spiking Dataset) and deep-neural-network-layer identification. By incorporating neural population preferences into model evaluation, framed RSA enables more mechanistically meaningful model comparisons and benefits from improved power for model-comparative inference.

**Significance Statement:** How should we compare representations across different systems, such as brain regions and layers of artificial neural networks? A popular method, representational similarity analysis (RSA), captures the geometry of a region’s response patterns but discards its mean response preferences (e.g., stronger overall responses to faces than to places)—a powerful clue to computational function. We introduce framed RSA, which enriches RSA with these mean response preferences, along with the global location and orientation of the response patterns. Framed RSA unites the two major traditions of brain-activity analysis—regional activation and multivariate pattern geometry—in a single functional fingerprint. Across human fMRI, macaque neural recordings, and deep networks, this fingerprint improves the power to distinguish competing models of brain computation.

## Introduction

Biological and artificial neural networks represent the world through distributed population codes: activity patterns across the population of biological neurons or model units encode the content of perceptual or cognitive representations. Each activity pattern representing a stimulus or piece of cognitive content can be conceptualized as a point in the multivariate response space of a brain region or model layer. For a sufficiently rich set of stimuli, the ensemble of points characterizes both the tuning of each neuron (columns of the stimulus-by-response matrix) and the representational geometry of the response patterns (rows of the stimulus-by-response matrix).^12,14^ A central concern in cognitive neuroscience, computational neuroscience, and AI is how best to measure the degree of alignment of these population codes across different brain regions or deep neural network (DNN) layers.^24,27,28^ A suitable measure of representational alignment is essential for comparisons between brain representations within^25^ and across species,^15^ and for evaluating DNNs as models of brain computation.^8,11,29,30^ When choosing a measure of representational alignment, we must consider that the systems to be compared often differ in many ways. Some of these differences (such as which neuron implements which tuning function) may be idiosyncratic to the individual animal or model instance^18^ and inessential to the algorithm, while others (such as what information is explicitly represented) may be indicative of the algorithm implemented by each system.^12,14^ A key question, then, is which aspects of the neural code should inform the comparison of two systems.

One common approach to comparing brain regions or DNN layers is *representational similarity analysis* ^13^ (RSA). In RSA, the pairwise distances in the multivariate response space among the neural representations of different stimuli or conditions are estimated and assembled in a square matrix, called the representational dissimilarity matrix (RDM). An RDM characterizes the *representational geometry* of a brain region or model layer. RDMs are then compared between different systems. In neuroscience, researchers often use correlation coefficients (e.g., Pearson’s *ρ* or Spearman’s *r*) between RDMs to measure representational alignment.^13,19,23^ In machine learning the linear centered kernel alignment is the most widely used comparator.^10^ These approaches are closely related,^5,27^ characterizing the representational geometry using distance (or more generally dissimilarity) or kernel (similarity) matrices, respectively. Focusing representational comparisons on the neural-population representational geometry, as these methods do, enables abstraction from the tuning of individual units and obviates the need for estimating an explicit linear or one-to-one mapping between two representations. Representational geometry provides a computationally informative abstraction in that it captures both the information content (assuming the noise is isotropic or has been isotropized) and the format in which that content is represented (up to an orthogonal linear transform).^12,14^ In particular, the representational geometry determines the accuracy with which any continuous or categorical variable can be linearly read out by a downstream decoder with access to all units in the population.

That said, comparing systems based on their representational geometry disregards other aspects of the neural population code, some of which may be biologically or computationally important. Notably, RSA is invariant to the global location and orientation of the ensemble of response patterns in the multivariate response space, since globally translating or rotating the set of patterns leaves their pairwise distances, and thus their geometry, unchanged. However, responses are nonnegative in both real neurons and DNN units (for common choices of the activation function). Biological neurons additionally have a limit to their maximum firing rate. The population response space of biological neurons is thus confined within a “hypercube” frame, a constraint which is ignored by standard RSA. The location and orientation of patterns relative to this frame often reflects computationally or biologically important aspects of the neural code, such as the tuning of individual neurons (e.g., whether they are tuned to a single experimental variable or exhibit mixed selectivity^22^ to multiple such variables), which in turn determines the neural wiring patterns required to encode and decode particular content. The global orientation and location of the response-pattern ensemble also affect the regional population-mean responses for a set of stimuli (such as a higher overall activation to images of faces than to images of houses in the fusiform face area). In particular, certain rotations can invert the population-mean response preferences of a region while leaving the distances between patterns unchanged^21^ (Fig. 1).

**Figure 1:**
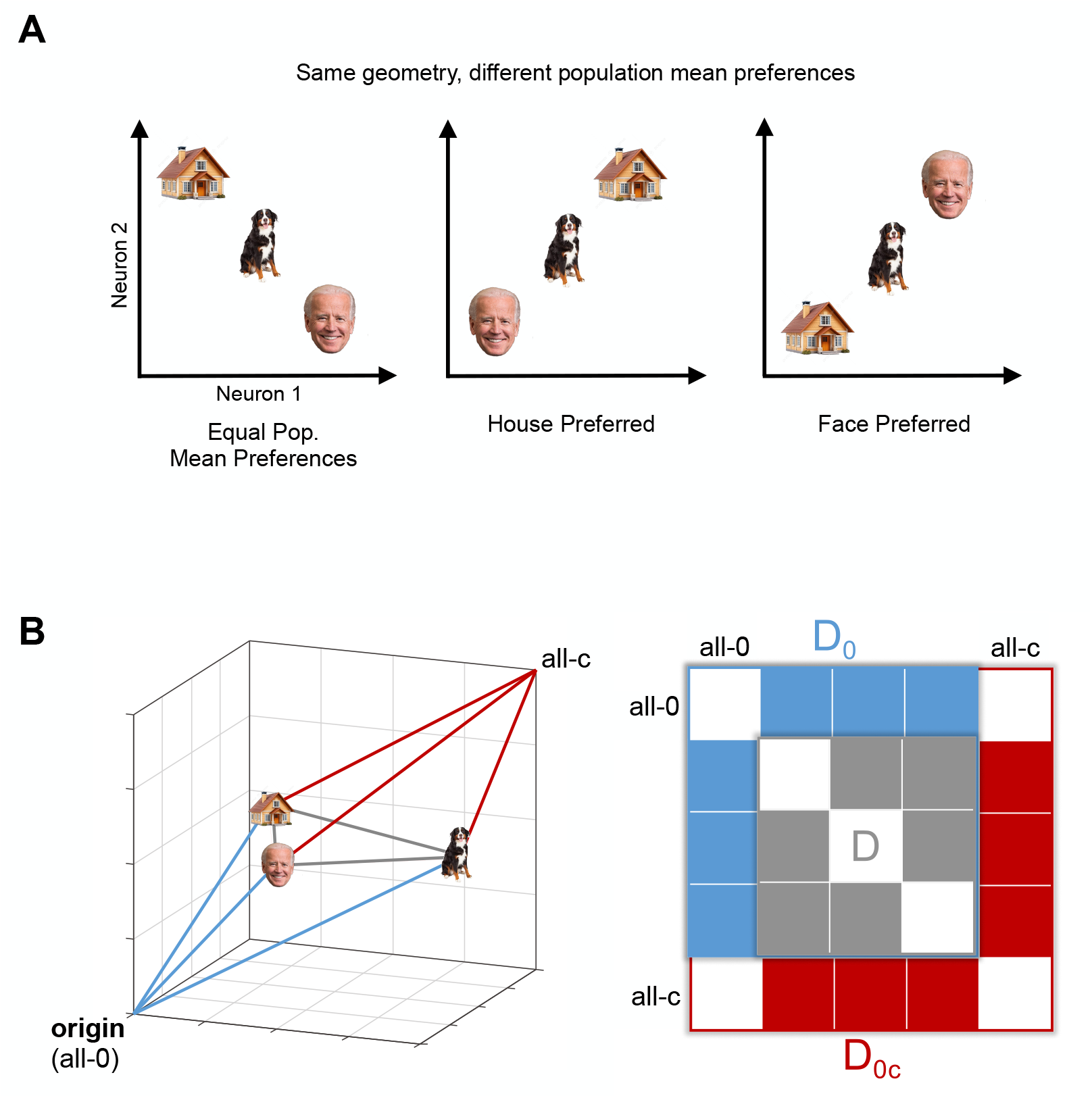
Motivation and schematic illustration of framed RSA. **(A)** Representations can have the same geometry (three panels), but distinct profiles across stimuli of the population-mean response. In the left panel, all stimuli elicit the same population-mean response. In the middle panel, the house elicits the strongest population-mean response. In the right panel, the face elicits the strongest population-mean response. Two brain regions that expend most of their energy on stimuli of different categories likely serve different computational functions. **(B)** By adding the all-zero (origin) and all-*c* patterns to the geometry, framed RSA creates a “reference frame” that renders RSA sensitive to different population-mean response profiles (i.e. to the differences among the three representations shown in A). Framed RSA removes the translational invariance and slightly reduces the rotational invariance that characterizes standard RSA. The distances from the stimuli to the origin (all-zero, blue) can be added to the initial RDM **D** to form the extended RDM **D**_**0**_, and the distances from the stimulus patterns to the all-*c* (red) can be further added to form the framed-RSA RDM **D**_**0c**_.

The regional population-mean response is a biologically important quantity in that it determines the metabolic cost of representing each stimulus. Augmenting RSA with information about the global location and orientation of stimulus patterns promises to capture biologically and computationally important facets of the neural code to which it is otherwise blind, and thus to enable it to better distinguish models that make different predictions about these aspects of the code. By contrast, other invariances of standard RSA may be worth preserving. For example, RSA is invariant to the number of units in a system (so long as there are enough units to preserve all pairwise distances), which is often desirable since the representations being compared nearly always differ in dimensionality. RSA is also invariant to the global scaling of the patterns because typical RDM comparators, such as correlation coefficients and the cosine similarity, are unaffected when the response patterns in one or both of the representations are all multiplied by a scalar constant. This invariance to joint scaling of all patterns is desirable for comparing systems with incommensurate activity units (e.g., spike rates in single-neuron recordings, BOLD responses of voxels, and activation values of DNN units).

Here we introduce *framed RSA*, a new variant of RSA that incorporates information about the global location and orientation of the pattern ensemble relative to the all-1 vector in multivariate response space, which renders RSA sensitive to the regional population-mean response preferences. This is achieved by augmenting the set of stimulus patterns with two “framing patterns”: the all-zero pattern and an all-*c* pattern, where *c* is a positive real number that defines the size of the multivariate-response hypercube (Fig. 1). For each stimulus, the distances between its activity pattern and these two framing patterns determine the population-mean response. As a consequence, framed RSA has reduced rotational invariance: it is not invariant to rotations that change the population-mean response for any of the stimuli. This extension integrates the traditionally separate cognitive neuroscience analysis of regional-mean activation across experimental conditions^2,6^ into RSA, combining representational geometry and the profile of mean activations into a single functional fingerprint for a brain region.

We first describe the mathematics underlying framed RSA, as well as how it interacts with various methodological choices in RSA. We then compare the performance of alternative variants of RSA at brain-region and DNN-layer identification, where ground truth is known, in order to validate framed RSA and explore different methodological choices. We show that framed RSA offers substantially improved power for model-comparative inference over both traditional RSA and mean-activation profile comparison. Finally, we make recommendations for best practices when using framed RSA. Framed RSA has been incorporated into the open-source RSA toolbox,^23^ making it easy to integrate into existing analysis pipelines.

## Results

### Mathematical overview

In framed RSA (Fig. 1), we augment the set of patterns defining the geometry with two additional patterns: the origin or all-zeros vector **0**, and a vector **c** consisting of some fixed positive constant *c*:

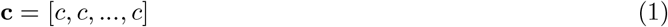

These two patterns define a hypercube in multivariate response space that constitutes a *reference frame* for determining the global location and orientation of the stimulus patterns. To aid intuition and emphasize the special role of these patterns, in visualizing the resulting RDM we set **0** as the first pattern and **c** as the last pattern, such that the entries corresponding to the distances to these patterns form a square “frame” around the RDM, further motivating the name of the method. We use **D** to denote the RDM containing the stimulus patterns alone, **D**_**0**_ for the RDM with the stimulus patterns and **0**, and **D**_**0c**_ for the RDM containing the stimulus patterns, **0**, and **c**.

Introducing **0** and **c** to the geometry endows the RDM with additional information about the location and orientation of the stimulus patterns, removing several forms of invariance present in standard RSA. The RDM **D**_**0c**_ is sensitive to global translations of the response-pattern ensemble since these can change the distances to **0** and **c**. The RDM **D**_**0c**_ is also sensitive to rotations of the stimulus patterns that change the distances to **0** and **c**. Note that rotations about **c** do not change the distances to **0** and **c** and so framed RSA remains invariant to these rotations.

While other choices of reference patterns would also provide information about the location and orientation of the stimulus patterns, the positions of patterns relative to **0** and **c** are of special computational relevance for several reasons. First, distances to **0** uniquely correspond to the norms of the patterns. Specifying these norms renders the RDM **D**_**0**_ informationally equivalent to the kernel matrix, such that these matrices are related by an invertible linear transform (see Methods). Second, the hypercube defined by **0** and **c** bounds the positive orthant of multivariate response space, i.e., the orthant in which all responses are positive. This orthant has a privileged status for biological and artificial neural networks, since firing rates in biological neurons and in artificial neurons with ReLU or logistic activation functions are necessarily non-negative. Other activation functions in artificial neural networks can yield negative responses, but most of them still compress negative but not positive responses (e.g., Swish/SiLU, Leaky ReLU, PReLU, ELU, SELU). Including the distances to **0** and **c** in the RDM does not fully define the location of the pattern ensemble, but it defines the eccentricity of the pattern ensemble within the positive orthant, i.e., the distance of the centroid from the all-*c* axis.

Finally, and crucially, including the distances to **0** and **c** in the RDM allows the regional population-mean response elicited by each stimulus to be recovered via linear operations on RDM entries when Euclidean distance (or certain variants thereof, such as Mahalanobis^26^ or crossnobis^5^ distance) is used as the dissimilarity measure. This can be shown by expanding the formula for the squared Euclidean distances among **0, c**, and a stimulus pattern **s**:

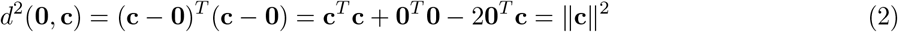

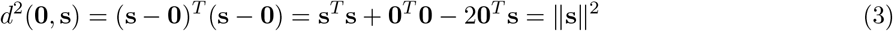

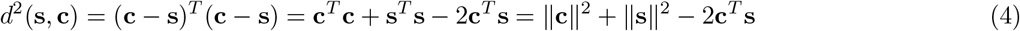

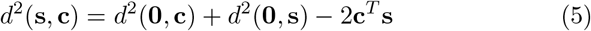

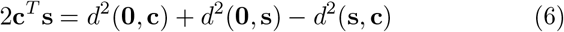

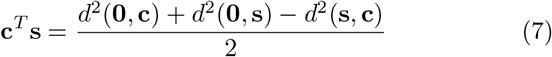

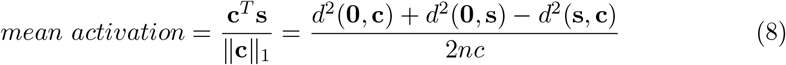

The population-mean activation is thus the inner product **c**^*T*^ **s** divided by a constant factor determined by the value of the fixed constant *c* and the number *n* of neurons or voxels.

#### Choosing the value of *c*

How should the value of *c* be chosen? It is evident from inspection that the overall geometry will change based on the value of *c*, since changing *c* will change all *d*^2^(**s, c**) entries while keeping all between-stimulus distances *d*^2^(**s**_*i*_, **s**_*j*_) the same. In particular, it can be geometrically demonstrated that for two values **c**_1_ and **c**_2_, the (vectorized) RDMs 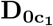 and 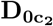 will be interconvertible via an affine (though not a linear) transform.

At minimum, methods for choosing *c* ought to obey the following constraints, motivated by preserving desirable invariances present in standard RSA. First, we wish for framed RSA to remain invariant to uniform multiplication of all channels by a fixed positive constant, since it is undesirable for an arbitrary change of measurement units to affect the results (e.g., converting raw BOLD activation to percent signal change), and since there is often no principled conversion factor when comparing different systems (e.g., neuronal firing rates and DNN unit activations). Second, the procedure for choosing *c* should not rule out the possibility of two systems having different numbers of units, but the same **D**_**0c**_, so long as all inter-stimulus distances are the same (or vary only by a fixed constant) between systems, and so long as all ratios between regional mean activations for different stimuli are the same.

Based on these considerations, we adopted the approach of choosing the constant *c* such that the resulting vector **c** has norm equal to the mean norm of the stimulus vectors times a constant *k*. This procedure guarantees that **D**_**0c**_ will remain unchanged by an arbitrary rescaling of measurement units, since *c* will be rescaled by the same factor. Moreover, it allows for systems with different numbers of units to have the same **D**_**0c**_ matrices. To see this, imagine taking a set of stimulus patterns in 2D along with the **0** and **c** vectors, and embedding the ensemble of patterns in 3D, such that the geometry is unchanged (residing in a 2D linear subspace), the **0** pattern is mapped to the origin in 3D space, and the **c** pattern falls on the equiangular axis (i.e., consisting of all constant-value vectors). The resulting 3D geometry will then have the same **D**_**0c**_ as the original 2D geometry. However, the value of *c* must be adjusted for the norm of **c** to match between 2D and 3D: 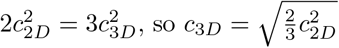. Choosing *c* such that the norm of **c** matches the mean norm of the patterns implies that while the *value* of *c* may vary across the systems being compared, the ratio *k* of its norm to the mean norm of the other patterns ought to remain constant.

The remaining free parameter then becomes the choice of *k*. As a pragmatic default, we let *k* = 1, such that **c** has mean norm equal to the other patterns. However, in the region identification analyses presented later we also explore how results depend on the value of *k*.

We note that tuning the value of *c* based on the mean norm of the stimulus patterns is not the only approach meeting the constraints described above; for instance, tuning *c* based on the mean projection of the stimulus vectors onto **c** would also preserve the desirable invariances described above.

#### Pattern-error whitening: Using noise-normalized dissimilarity estimators in framed RSA

To account for varying levels of noise in different channels and for correlated noise between channels, it is often desirable to estimate the Mahalanobis distance. We can estimate the Mahalanobis distance either by naively computing it from the data (where the noise inflates our estimate of the noisefree underlying distance, producing a positive bias) or using the *crossnobis* estimator,^19,23,26^ which removes the bias by crossvalidation:

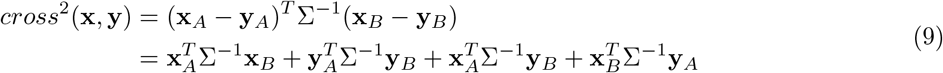

where *A* and *B* refer to different data partitions (e.g., fMRI runs); if A=B this reduces to standard Mahalanobis distance. In the context of framed RSA, if we follow the same steps as for Euclidean distance, we can use *cross*^2^(**0, c**), *cross*^2^(**0, s**), and *cross*^2^(**s, c**) to solve for **c**^*T*^ Σ^−1^**s**, which reflects not the raw regional mean as with Euclidean distance, but rather the *noise-normalized* regional mean. If only univariate noise normalization is applied, this corresponds to rescaling each channel by its noise variance before computing the mean. If multivariate noise normalization is applied, this corresponds to first rescaling the activations based on the principal components of the noise covariance prior to taking the mean (i.e., accounting for correlated noise between channels).

Mahalanobis and crossnobis distance raise an additional complication when it comes to choosing the value of *c*: should the norm of **c** be chosen based on the mean norm of the stimulus patterns before or after whitening? Consider two systems whose geometries are distinct before whitening, but become identical after whitening. Applying standard RSA with Mahalanbis/crossnobis as the dissimilarity metric would regard these systems as the same; if we wish for the same to be true for framed RSA, the norm of **c** must be tuned based on the norms of the patterns after whitening. This is therefore the method we adopt.

#### RDM-error whitening: Using whitened RDM comparators in framed RSA

A simple way of comparing RDMs is to use functions such as cosine or correlation, which have the desirable properties of being invariant to arbitrary rescaling of measurement units. However, these have the disadvantage that they do not account for the covariance structure of the dissimilarity errors of the RDM.^5^ For instance, the distance estimates *d*^2^(**x, y**) and *d*^2^(**x, z**) share the same pattern estimate **x** whose error affects both of them. The dissimilarity errors between the RDM entries associated with these two distances are therefore correlated. Whitened variants of correlation and cosine can be used to account for this covariance structure, which has been shown to improve model-comparative power in some cases.^23^ These whitened variants require estimating and inverting **V**, the variance-covariance matrix of the dissimilarity values. **V** has a row and column for each off-diagonal entry of the RDM, with its diagonal entries containing the variance of each dissimilarity in the RDM, and its off-diagonal entries containing the covariance between all pairs of dissimilarities in the RDM. For instance, the entry of **V** corresponding to the covariance between *d*^2^(**w, x**) and *d*^2^(**y, z**) would be zero if all four patterns correspond to distinct stimuli with independent noise, since these two distances share no noise in common.

In past work, **V** has typically been estimated under the simplifying assumption that the true distances between patterns are zero.^5^ However, this approach is unsuitable for framed RSA for two reasons. First, the **c** and **0** vectors used in framed RSA are both noiseless; if we further assume the distance between them to be zero (such that neither signal nor noise distinguish them), their associated rows in **V** will be identical, resulting in a singular matrix. Second, since framed RSA involves choosing a scaling factor for **c**, it is desirable for **V** to take into account the magnitude of this scaling factor, which is impossible if we compute **V** under the assumption that the true distances are zero. That said, using noisy estimates of the underlying distances to compute **V** raises numerical challenges, since the size of **V** scales with the fourth power of the number of patterns (the number of pairs of pairs of stimuli), and imprecise estimates of the underlying distances may render the inversion of **V** numerically unstable. For these reasons, we therefore make several adjustments to the use of whitened RDM comparators in the case of framed RSA, as described in the Methods.

As mentioned earlier, for two different values of *c* ∈ {*c*_1_, *c*_2_}, the RDMs 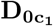 and 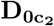 will be interconvertible via a transform that is affine, but not linear. Since whitened cosine and whitened correlation are invariant to linear transformations of their inputs, but not to general affine transformations, RDM similarity will still depend on the value of *c* even when using whitened RDM comparators.

### Framed RSA exhibits superior power in region identification with fMRI and neural-recording data

A region identification paradigm was used as a proxy task to evaluate how powerfully framed RSA could adjudicate between different computational models. Since framed RSA includes information about both the representational geometry and regional mean activations of a neural population, its performance was also compared to standard RSA and to the profile of mean activations, in order to characterize how it might perform relative to either kind of information in isolation. This approach was applied across two neural datasets.

#### Natural Scenes Dataset

Region identification results for the Natural Scenes Dataset are shown in Fig. 2. Framed RSA significantly outperforms standard RSA at all stimulus sample sizes when whitened correlation or whitened cosine similarity were used as RDM comparators, with weaker or mixed results when non-whitened correlation or cosine were used as RDM comparators. Framed RSA outperformed the profile of mean activations for small numbers of stimuli, but was significantly outperformed at higher numbers of stimuli.

**Figure 2:**
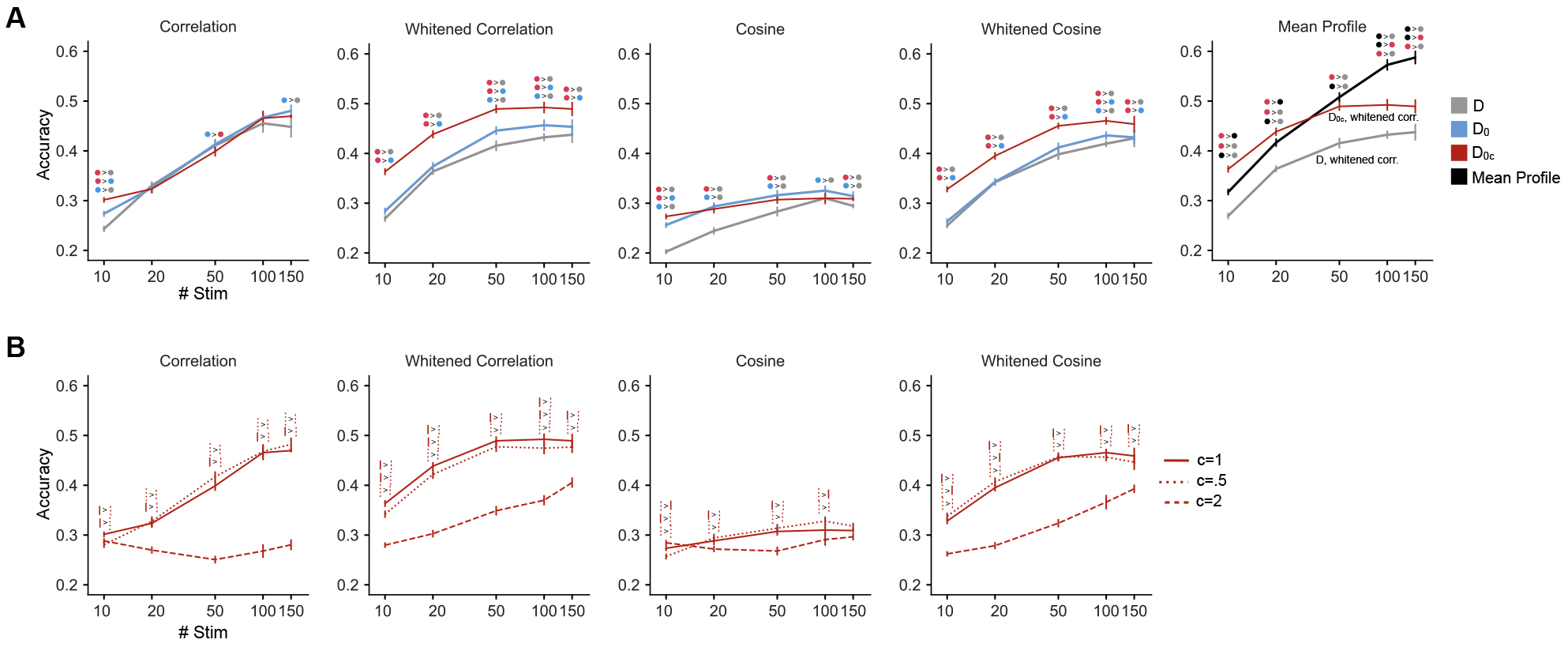
Region identification results for the Natural Scenes Dataset. **(A)** Region identification accuracy for framed RSA (red, summary statistic: **D**_0*c*_), framed RSA with **c** omitted (blue, summary statistic: **D**_0_), standard RSA (gray, summary statistic: **D**), and profile of mean activations (black) for different RDM comparators. Significant pairwise contrasts (matched-pairs *t*-tests; *p <* .05) are denoted using the icons above each datapoint. To compare RSA variants to the profile of mean activations, the results for Framed and Standard RSA with their highest-performing comparator are re-plotted on the mean profile panel. **(B)** Region identification performance results when the length of **c** relative to the mean length of the stimulus vectors is varied.

Next, we examined the effects of varying the ratio of the norm of *c* relative to the mean norm of the stimulus vectors. We found that a ratio of 0.5 or 1 outperformed a ratio of 2 across all RDM comparators, with little difference between 0.5 or 1.

#### Things Ventral Stream Spiking Dataset

Region identification results for the Things Ventral Stream Spiking Dataset are shown in Fig. 3. Framed RSA yields higher region identification performance than standard RSA at low numbers of stimuli across all RDM comparators. For high numbers of stimuli, framed RSA performs similarly to standard RSA when unwhitened correlation or cosine are used as RDM comparators, but lower performance when their whitened variants are used. Framed RSA outperforms the profile of mean activations for all numbers of stimuli.

**Figure 3:**
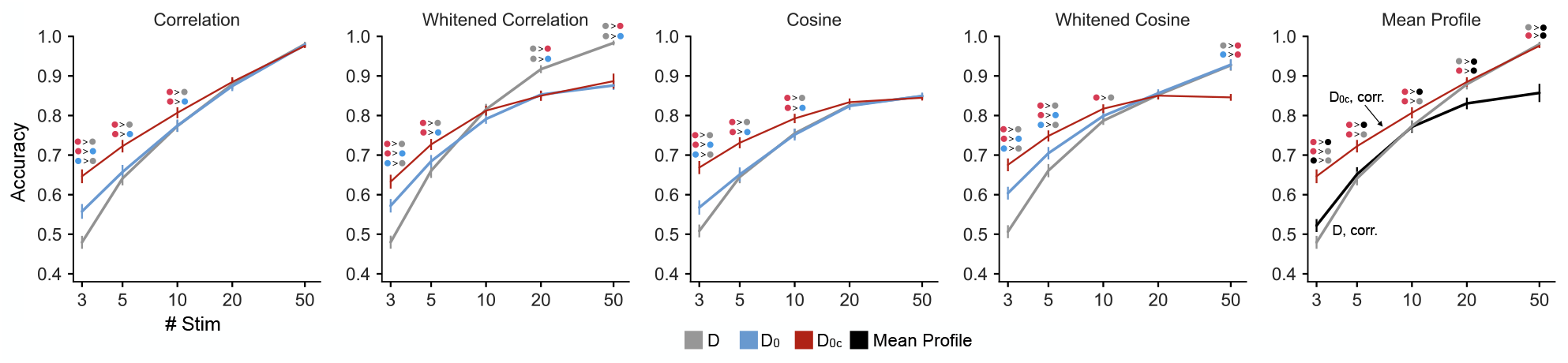
Region identification results for the TVSD. Significant pairwise comparisons (matched pairs *t*-tests, *p <* .05) are denoted with icons above each datapoint. To compare the accuracy for the profile of mean activations to standard and framed RSA, these analyses are re-plotted on the Mean Profile panel, using the RDM comparator that yielded the highest overall accuracy.

When using the best-performing RDM comparator for each method (correlation for framed RSA, whitened correlation for standard RSA), framed RSA thus matches or exceeds region identification performance of alternative methods.

### Framed RSA exhibits superior power in DNN layer identification

We then extended this region identification approach to DNNs to characterize how well framed RSA could correctly match corresponding DNN layers across different random seeds compared to other alignment metrics. DNN layer identification results are shown in Fig. 4, at low (**A**) and high (**B**) levels of noise. At low levels of noise, framed RSA significantly outperforms standard RSA and the profile of mean activations at all numbers of stimuli, and for all RDM comparators. At high levels of noise, framed RSA substantially outperforms standard RSA across all comparators and numbers of stimuli. However, while it outperforms the profile of mean activations when few stimuli are used, at higher numbers of stimuli the profile of mean activations yields higher accuracy. Both representational geometry and mean activation information therefore seem to independently contribute to differentiating between layers of AlexNet in a way that generalizes across random seeds, with mean activation information being especially helpful at higher levels of noise.

**Figure 4:**
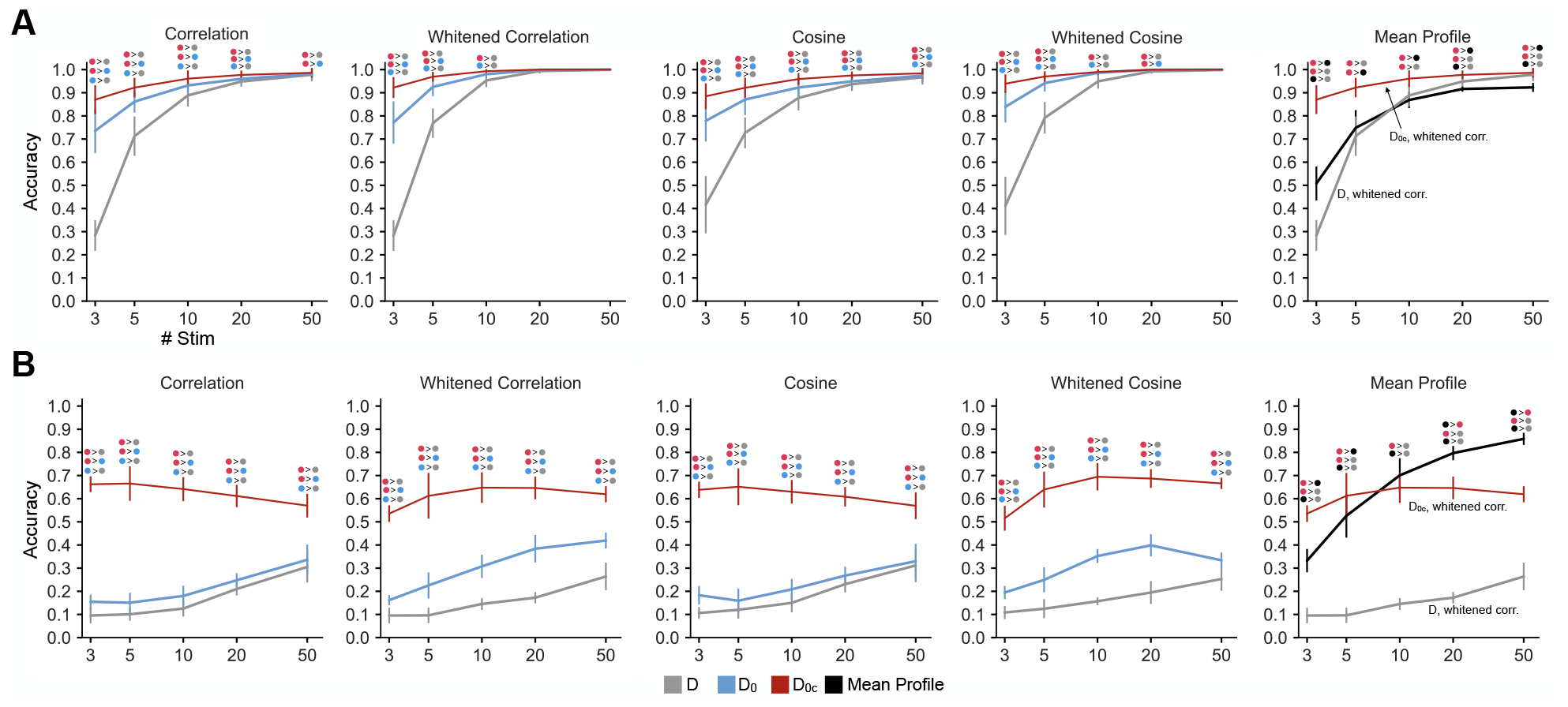
DNN layer identification performance results. Layer identification accuracy is shown for low (A) and high (B) levels of noise. Significant pairwise comparisons (matched pairs *t*-tests, *p <* .05) are denoted with the icons above each datapoint. To compare the accuracy for the profile of mean activations to standard and framed RSA, these analyses are re-plotted on the Mean Profile panel, using the RDM comparator that yielded the highest overall accuracy.

## Discussion

Comparing neural-population representations requires us to decide what aspects of the representations we consider signatures of computational function, and what aspects we consider nuisance variation. Here we introduce framed RSA, which combines information about both the representational geometry and the regional mean stimulus preferences of a neural population through the simple approach of adding the origin and a constant pattern to the geometry as additional reference points in the multivariate response space, thus providing a reference frame that defines the global location and orientation of the stimulus patterns relative to the all-1 vector.

To objectively assess the model-adjudication performance of framed RSA and to compare its performance to standard RSA and to matching up regions according to the similarity of their regional-mean activation profiles, we used model-recovery simulations, where ground truth is known. In particular, we used both brain-region-identification analyses (across individual humans whose brain activity was measured with fMRI and across macaque monkeys undergoing neural array recordings), and DNN layer-identification (across instances of DNN models).

For the Natural Scenes fMRI dataset, framed RSA either matched or outperformed standard RSA across all RDM comparators or numbers of stimuli. Framed RSA also significantly outperformed matching by regional-mean activation profile for small numbers of stimuli (≤ 20). Interestingly, however, when many stimuli (≥ 100) were used, the profile of mean activations performed best of all. One explanation for these results is that the regional mean is robust to noise, since averaging across voxels will tend to cancel out the effects of noise. For small numbers of stimuli, the regional-mean activation profile may not be rich enough to provide a unique signature for identifying regions with fMRI. For large numbers of stimuli, however, the regional-mean activation signature appears to be rich enough for identifying an ROI among the regions tested here, which included retinotopic regions V1-hV4 in addition to category-selective regions defined based on their sensitivity to faces, bodies, scenes, or words (for which the profile of mean activations may be an especially informative functional signature).

Interestingly, for the NSD dataset we observed that region identification accuracy was higher when correlation was used as an RDM comparator than when cosine was used, for both the whitened and unwhitened variants of these comparators, and across all versions of RSA. Cosine similarity is sensitive to uniform shifts in the RDMs being compared (i.e., adding a constant offset to all RDM entries), such that there is a meaningful zero point and ratios between distances are retained;^5^ by contrast, correlation similarity is invariant to uniform shifts in the RDM, discarding the ratios between distances. In the context of the region identification paradigm, the magnitude of the constant RDM offset thus does not seem useful for distinguishing brain regions in a manner that generalizes across subjects.

The fMRI region-identification analyses also showed the choice of *c* matters to the performance of framed RSA: making *c* too large greatly reduced performance across various RDM comparators, regardless of whether RDM whitening was applied. A potential explanation for this observation may be found in the decomposition for *d*^2^(**s, c**) given in Equation 4: as *c* increases, the ∥**c**∥^2^ term increasingly dominates the overall sum for *d*^2^(**s, c**), while leaving the stimulus-to-stimulus dissimilarities unaffected, progressively widening the gap between the *d*^2^(**s, c**) RDM entries and stimulus-to-stimulus RDM entries (i.e., such that the vectorized RDM increasingly resembles a step function). At the same time, increasing *c* also scales up the **c**^*T*^ **s** component of *d*^2^(**s, c**), magnifying the noise in these entries. The joint consequence of these effects is that increasing **c** may cause increasingly noisy entries to dominate RDM comparison, potentially impairing region identification performance.

For the macaque neural-recording dataset (TVSD), framed RSA always beat or matched standard RSA in performance when each was used with its best-performing RDM comparator. The best-performing RDM comparator was Pearson correlation for framed RSA and whitened Pearson correlation for standard RSA. Framed RSA significantly outperformed standard RSA only for small numbers of stimuli (≤ 10), but outperformed the profile of mean activations at all numbers of stimuli. Given the lower levels of noise in neural recording data relative to fMRI, it is possible that the regional-mean activation exploited by framed RSA adds little extra information relative to the fairly fine-grained information already present in the neural population patterns. For the DNN dataset, framed RSA outperformed standard RSA at both low and high levels of noise, and outperformed the profile of regional-mean activations at low levels of noise, but not at high levels of noise. These results corroborate the interpretation that the regional-mean activation profile may be especially diagnostic at higher levels of noise. In the context of DNNs, framed RSA may be especially useful for distinguishing ReLU layers from other layers, since their activations are necessarily confined to the positive orthant of response space where the all-*c* pattern resides.

Framed RSA is less rotationally invariant than standard RSA, but just how sensitive to rotation is it? The number of degrees of rotational freedom is the number of pairwise orthogonal planes of rotation. In an *n*-dimensional space, the number of rotational degrees of freedom is therefore the number of pairs among the *n* orthogonal axes: 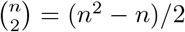. For three neurons, for example, the number of rotational degrees of freedom is 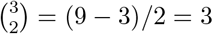. Framed RSA removes one dimension, leaving 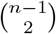 degrees of rotational freedom. For three neurons, this leaves only 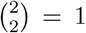 degrees of rotational freedom (rotations about **c**). However, for a population of *n* ≫ 3 units, the number 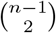 of degrees of rotational freedom—that is, the number of ways the geometry can rotate without changing **D**_**0c**_—is large. There are *n* − 1 distinct axes orthogonal to **c**, with any given pair of these axes defining a unique plane of rotation that leaves **D**_**0c**_ unaffected. For *n* = 100 units, for example, there are 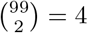, 851 remaining degrees of rotational invariance for framed RSA, whereas in conventional RSA there are 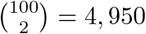. In general, framed RSA reduces the number of rotational degrees of freedom by factor 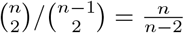. This amounts to only a slight reduction of rotational invariance when the dimensionality is high.

Existing measures for brain-model alignment fall on a spectrum from rigid to flexible [28]. More rigid methods, such as one-to-one matching, have smaller equivalence classes of representations because they have stricter requirements for considering two representations similar. More flexible methods, such as linear encoding models, have large equivalence classes because they have looser requirements for considering two representations similar. RSA occupies a middle ground on this spectrum, disregarding rotations, but requiring representations to place equal relative emphases on all representational dimensions to be considered equivalent. Framed RSA extends the RSA framework toward somewhat greater rigidity, by additionally requiring regional-mean activation profiles to be positively correlated. Framed RSA is also additionally sensitive to translations of the response-pattern ensemble. However, framed RSA is much less rigid than soft-matching,^9^ a generalization of one-to-one matching of tuning functions to systems with different numbers of neurons (or measured response channels).

In conclusion, framed RSA offers a simple approach for incorporating both representational geometry and regional-mean information into model comparison, offering model-brain comparisons that honor the regional-mean response preferences of brain regions. The population-mean response reflects the energy required by a region to represent or process different stimuli and is therefore an important functional signature of brain computation that models ought to correctly predict. Requiring models to predict this feature along with the geometry also promises more powerful model comparison.

## Methods

### Relationship between Euclidean distance RDM and the kernel matrix

The kernel matrix **K** (also called the Gram matrix) is the matrix of inner products between all pairs of stimulus patterns, with the squared pattern norms along the diagonal. Here, we demonstrate that each entry of **K** can be recovered via linear operations on the entries of **D**_**0**_ (the RDM of squared Euclidean distances among all stimulus patterns and the all-zeros vector **0**) and vice-versa. As a result, the kernel matrix **K** and the RDM **D**_**0**_ are related by an invertible linear transform and thus informationally equivalent.

Each entry of **D**_**0**_ is the squared Euclidean distance between two patterns **x** and **y**, which can be written as:

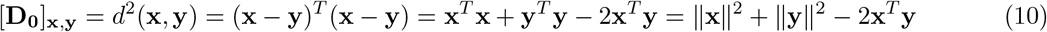

Since ∥**x**∥^2^, ∥**y**∥^2^, and **x**^*T*^ **y** are entries in **K**, every entry of **D**_**0**_ can be recovered via linear operations on **K**.

In the reverse direction, any entry **x**^*T*^ **y** of **K** can be recovered from entries of **D**_**0**_ via:

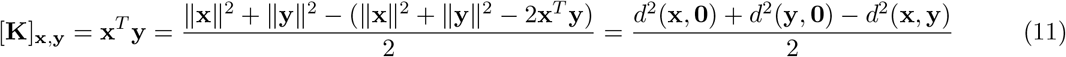

Thus, any entry of **K** can be recovered via linear operations on entries of **D**_**0**_ and vice-versa, such that either (flattened) matrix can be converted to the other via an invertible linear transform. However, the matrix **D** (i.e., without the distances to **0**) does not have this property, since it does not contain the squared pattern norms *d*^2^(**x, 0**) and *d*^2^(**y, 0**) needed in equation 11.

### Using whitened RDM comparators in framed RSA

Using whitened RDM comparators in framed RSA requires several mathematical adjustments relative to how these comparators are used in standard RSA. We describe these adjustments here.

Applying whitened RDM comparators, such as whitened cosine or whitened correlation, requires estimating the variance-covariance matrix of dissimilarity estimates **V**, whose entries are the variance of each off-diagonal entry in the corresponding RDM **D** and the covariance between each pair of off-diagonal entries. The values of **V** depend on the magnitudes of both the true underlying distances between patterns and of the noise.^3,4,5^ It is often convenient to estimate **V** under the assumption that the true underlying distances are zero,^5^ since noisy distance estimates can sometimes lead to numerical instability when **V** is inverted, potentially negating the advantages of using whitened comparators in the first place. However, in framed RSA taking the true distances into consideration when estimating **V** is essential, since otherwise **V** is non-invertible. The reason for this is as follows. Since the **0** and **c** patterns are noiseless, treating the distance between them as zero (i.e., as if they were the same pattern) makes them indistinguishable, such that for any stimulus pattern **s**, the rows of **V** corresponding to the distances *d*^2^(**s, 0**) and *d*^2^(**s, c**) will be identical (i.e., these distances will covary with the other distances in exactly the same way), making **V** singular.

A prior method for estimating **V** for the crossnobis distance makes the simplifying assumption that the true mean signal and the noise have the same spatial covariance structure.^4,5^ However, we can derive an improved formula for **V** that does not make this assumption as follows.

Suppose we measure the responses of *P* channels to *K* stimuli with prewhitened true vector representations **U**_*i*_ summarized in one matrix **U**, whose noise is drawn from a matrix-normal distribution with covariance over voxels **Σ**_*R*_ (i.e., the remaining voxel covariance after whitening, which will equal the identity matrix if whitening is fully successful) and covariance over stimuli/conditions **Σ**_*K*_ (reflecting the extent to which noise covaries across different conditions, e.g. due to temporally sustained BOLD noise affecting adjacent trials in an fMRI experiment).

We define the column vectors ***δ***_*ij*_ = **U**_*i*_ − **U**_*j*_ and estimate these differences with their measured counterparts 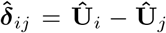 assuming that **Û** _*i*_ = **U**_*i*_ + ***ε***_*i*_, where ***ε***_*i*_ is drawn independently with zero mean across *M* separate crossvalidation folds (e.g., fMRI runs).

We want to look at the covariance of two Crossnobis distance estimates of the form

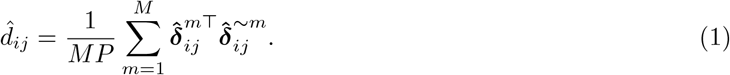

Here the superscript *m* refers to the estimate from the *m*-th cross-validation fold, and ~*m* refers to the mean estimate from the union of all other folds.

Linearity of the covariance leads to the following formula for the covariance of two distance estimates as in Diedrichsen et al.:^4^

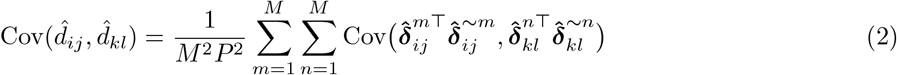

Now we want to calculate the individual covariance terms 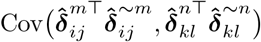 without the second two assumptions made by Diedrichsen et al.^4^ in their Equation 27, which stipulate that the spatial structure of the noise and the true mean signal are the same (i.e., both equal to **Σ**_*R*_ scaled by different constants).

To reduce clutter in the notation we follow Diedrichsen et al.^4^ and first expand 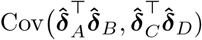 for general vectors 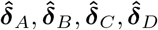 with 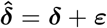 using only the information that (*A, B*) and (*C, D*) are from separate runs, i.e. ***ε***_*A*_ ⊥ ***ε***_*B*_, ***ε***_*C*_ ⊥ ***ε***_*D*_, and ***ε***_*A*_ ⊥ ***ε***_*C*_ | ***ε***_*A*_ ⊥ ***ε***_*D*_ and ***ε***_*B*_ ⊥ ***ε***_*C*_ | ***ε***_*B*_ ⊥ ***ε***_*D*_.

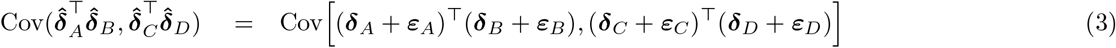

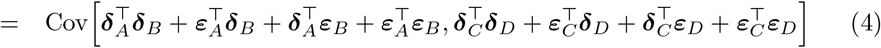

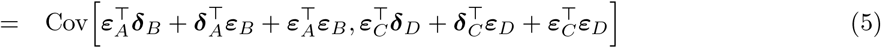

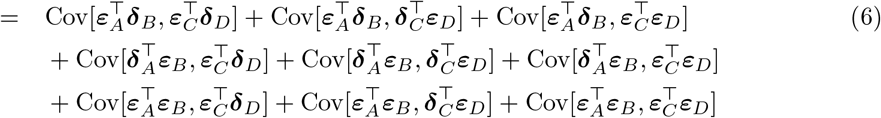

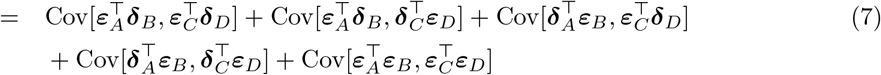

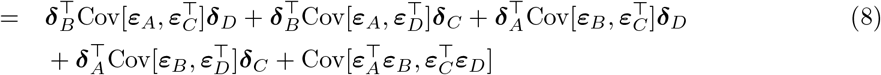

Now let us use the definitions for the error vectors:

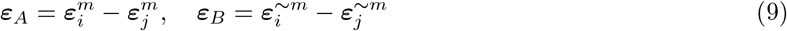

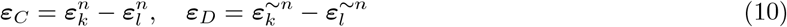

With these we calculate the five individual covariances from equation 8:

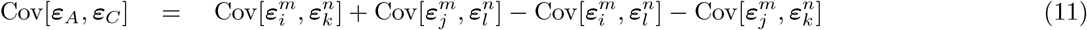

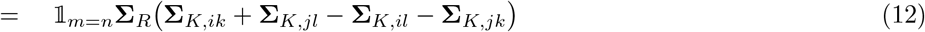

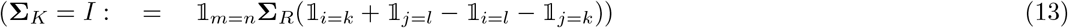

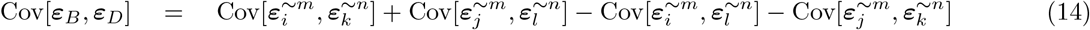

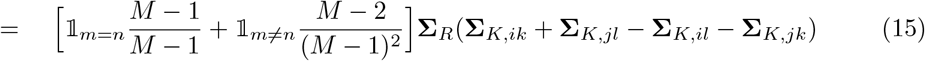

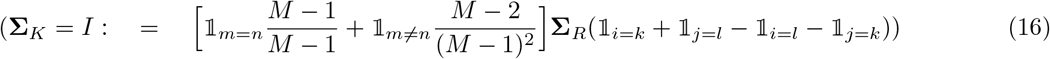

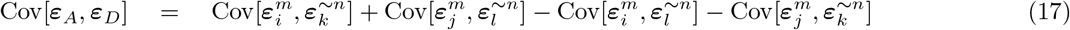

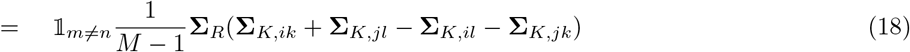

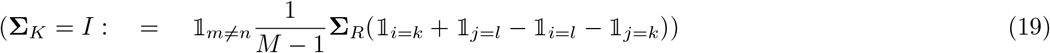

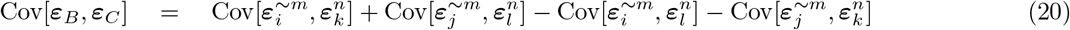

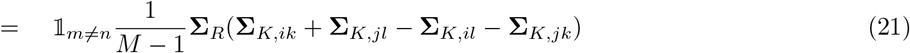

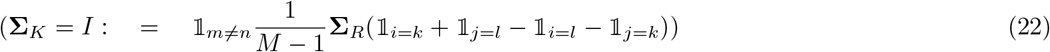

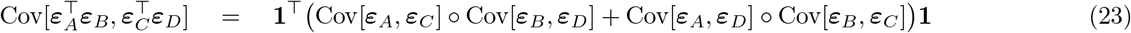

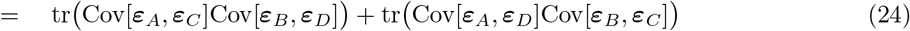

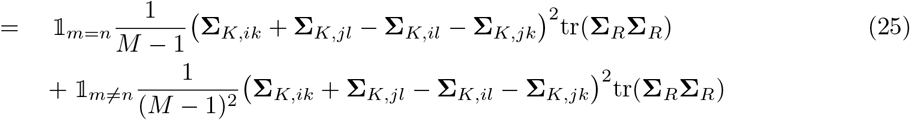

Now look at the full individual terms in equation 8, while defining 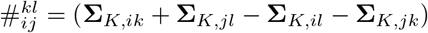:

If *m* = *n*:

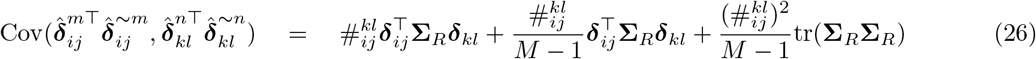

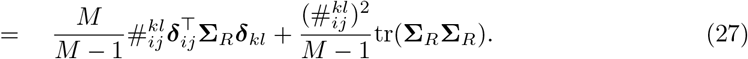

If *m*≠ *n*:

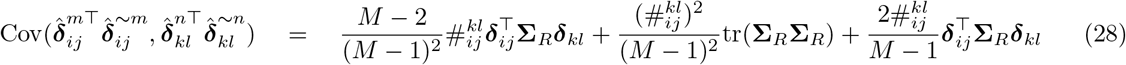

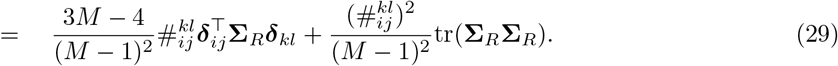

Finally, plugging these into equation 2:

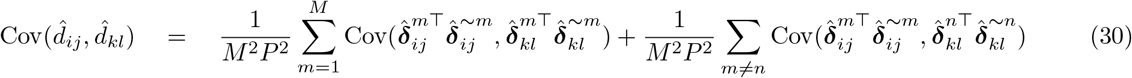

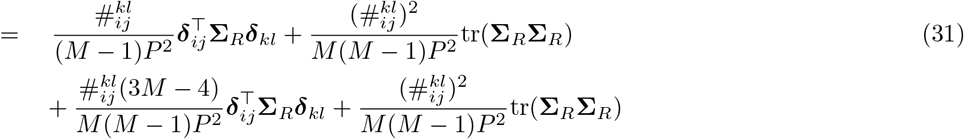

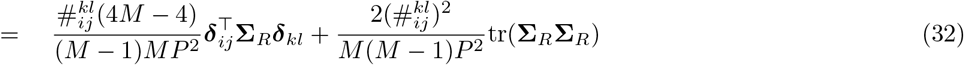

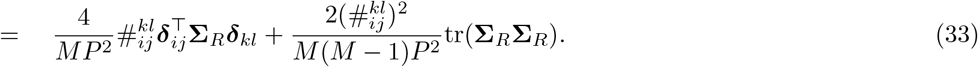

Which can be written in matrix form using the contrast matrix **C** and 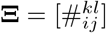 as:

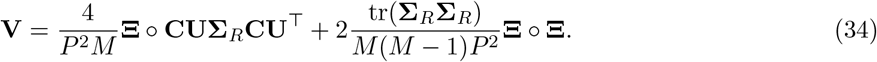

Thus, **V** is the sum of a signal-dependent term (the first term in the sum in Equation 34) and a signal-independent term (the second term in the sum). As mentioned earlier, this formula removes the assumption from earlier work^3,4,5^ that the signal and noise have the same spatial covariance structure. The formula derived in past work is:

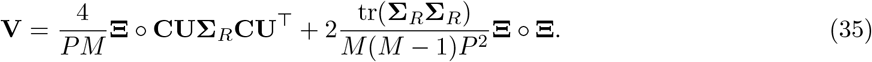

Thus, the signal-dependent part of the new formula differs from the corresponding term of the formula used in past work by a factor of *P*, whereas the signal-independent part is the same. That said, this formula can still yield numerically unstable estimates of **V**^−1^ in cases where the true patterns are not measured precisely, and in certain cases it may thus be desirable to only use the signal-independent term. Numerical instability when inverting **V** becomes especially problematic with large numbers of stimuli, since the number of entries in **V** scales with the fourth power of the number of stimuli (i.e., the number of pairs of pairs). One further point is that since **Σ**_*R*_ is the remaining noise covariance between channels that persists after whitening (since the true covariance will not be perfectly measured with limited data), when computing **V** for real data **Σ**_*R*_ will be treated as the identity matrix for practical purposes.

Using this formula in the context of framed RSA introduces several additional considerations. First, the rows and columns of **Σ**_*K*_ corresponding to **0** and **c** must be set to zero, since these patterns are noiseless. Second, the row and column of **V** corresponding to *d*^2^(**0, c**) must be removed (along with the *d*^2^(**0, c**) entry in the RDM), since they will consist of all zeros as a consequence of *d*^2^(**0, c**) being a constant value and having zero variance. Third, it is mandatory to use the signal-dependent term in Equation 34, at the very least for all entries of **V** involving **0** or **c**. The reason for this is that since these patterns are noiseless, if the distances between them are treated as zero then they will be fully interchangeable, such that rows/columns of **V** involving these patterns will be identical and render the matrix singular. By contrast, it is not essential to use the signal-dependent term for entries of **V** involving only the stimulus patterns, since their trial-specific noise is sufficient to individuate them.

Thus, we make the following practical recommendations when estimating **V** in the context of framed RSA. If the ground-truth stimulus patterns can be measured with sufficient precision, then it is appropriate to use Equation 34 as given, using both the signal-dependent and signal-independent terms for all entries of **V**. In cases where estimating and inverting **V** may be numerically unstable due to imprecise pattern estimates or a large number of stimuli, a practical approach is to set the rows of the contrast matrix **C** corresponding to distances between pairs of stimulus patterns to zero, keeping the other rows (i.e., distances between a stimulus pattern and either **0** or **c**) the same. This amounts to assuming that the stimulus-to-stimulus distances are zero for the purposes of estimating **V** (though not for the RDM itself, for obvious reasons), while still taking the framed-to-stimulus distances into consideration. We adopted this approach for the fMRI region identification analyses involving the Natural Scenes Dataset after observing that region identification accuracy surprisingly *decreased* with more stimuli if the full estimate of **V** (i.e., incorporating all estimated distances) was used to compare RDMs with whitened comparators, but substantially improved and increased monotonically with stimulus count if our adjusted procedure for estimating **V** (i.e., treating stimulus-to-stimulus distances as zero, while still incorporating the other measured distances) was used.

### Region Identification Experiments

Introducing **0** and **c** to the geometry allows the regional-mean activation evoked by each stimulus to be recovered as a linear function of the dissimilarities (Equation 8). The dissimilarities can be squared Euclidean, squared Mahalanobis, or crossnobis dissimilarity estimates for this linear relationship to the mean to hold (by contrast, other dissimilarity measures like correlation distance cannot be used to recover the regional mean). We will recover the raw population mean if squared Euclidean distance is used as the dissimilarity, or the noise-normalized mean if the squared Mahalanobis distance or the crossnobis estimator is used. The mean stimulus preferences of a region are of theoretical interest in their own right.

An important further question is whether this added information gives framed RSA more power to adjudicate among computational models of brain function relative to standard RSA. Evaluating model adjudication accuracy requires ground truth about the correct identification. We do not have the ground-truth model for any brain region (e.g., V1). However, we can use measurements performed in the same and other brain regions in other animals as proxies for the models to be evaluated. We then have ground truth: other animals’ V1 should be identified as the correct “model” of V1 in a given animal. We refer to this approach to validating alignment measures as *region identification*[17]. In region identification, we assess the effectiveness of a given method for model adjudication by determining how frequently it categorizes different instances of the *same* region (e.g., V1 of two different subjects) as more similar than instances of different regions (e.g., V1 in one subject and V2 in a different subject). For simplicity, we use the phrase “region identification” to encompass the application of this approach to DNNs as well, treating their layers as analogous to regions of the brain and specific model instances (same architecture and task trained from different random seeds) as analogous to different subjects.

We applied this approach across three datasets. The first dataset is the Natural Scenes Dataset^1^ (NSD), a massive open-source 7T fMRI dataset with responses to thousands of images from eight subjects, including responses to 1,000 shared images that were presented to all subjects. The second dataset is the Things Ventral Stream Spiking Dataset^20^ (TVSD), consisting of intracranial neural recording data from V1, V4, and IT of two macaque monkeys. The third dataset is a set of ten instances of AlexNet^16^ trained for image classification on ImageNet from different random seeds.

These datasets were chosen based on the following criteria. First, the region identification paradigm requires that multivariate pattern data can be collected from multiple clearly demarcated regions (layers in the case of DNNs) of multiple subjects (instances in the case of DNNs), with every region present in every subject. This requirement excludes modalities such as EEG or MEG, since these methods lack the spatial resolution to collect fine-grained response patterns from specific brain regions. The requirement that data from all regions be present in all subjects also limits the viability of using human intracranial data, since such data is typically collected opportunistically, with the location of electrodes and quantity of data being constrained by clinical factors. Second, the datasets are complementary in several useful respects. The NSD and TVSD involve different species and different measurement modalities with varying levels of noise, with fMRI being much noisier than intracranial recordings. Since the regional mean activation is plausibly robust to noise, it is of interest to examine how the potential benefits of framed RSA generalize across different noise levels. The inclusion of a DNN dataset offers the benefit of having access to the ground-truth stimulus patterns, and being able to simulate different levels of noise as desired.

The region identification analyses followed the same structure in humans, monkeys, and models, with adjustments for each dataset as needed. We now explain that shared structure, using the term “region” to encompass both brain regions and DNN layers, and the term “subject” to encompass both individual humans/monkeys and DNN instances. One of the subjects was designated as the test subject, and one of the regions as the true region. The data RDM was then computed for the true region in the test subject. An RDM was computed for all the regions (including the true region) in all other subjects. The RDMs for these subjects were then averaged to generate a single “model” RDM for each region. The similarity between the data RDM and each “model” RDM was computed, with a trial being tallied as “correct” if the data RDM was more similar to the “model” RDM from the true region than it was to the other “model” RDMs. The proportion of correct decisions was stored as the region-identification accuracy (RIA) for the data RDM. The RIA was computed for each combination of test subject and test region. The mean RIA across all test subjects and true regions was estimated for each measure of representational alignment and for different random draws of stimuli, varying the number of stimuli. The manner in which stimuli were sampled depended on the dataset, accounting for differences between the three datasets.

For all datasets, the measures of representational alignment we compared were (1) standard RSA using just the stimulus patterns (i.e., with the RDM **D**), (2) framed RSA including the stimulus patterns and **0** (**D**_**0**_), (3) framed RSA including the stimulus patterns, **0**, and **c** (**D**_**0c**_), and (4) the profile of mean activations. For all RSA methods, the crossnobis estimator was used as the dissimilarity, with univariate noise normalization. To examine how the benefits of framed RSA might depend on the choice of RDM comparator, (1)-(3) were run using correlation, whitened correlation, cosine, or whitened cosine to compare RDMs. For the whitened RDM comparators, an adjusted formula for **V**, the matrix of covariances among RDM entries, was used (as described in the previous section). For most analyses, **c** was set to have norm equal to the mean norm of the stimulus patterns, with the exception of an analysis in the NSD that compares RIA for different values of the norm of **c**. RIA was then statistically compared between representational alignment measures within each RDM comparator and number of stimuli (e.g., RIA for standard RSA vs. framed RSA when whitened correlation was used as the RDM comparator for draws of 10 stimuli at a time), with the exact statistical comparison method varying across datasets to account for differences in trial structure. The performance of standard RSA and framed RSA were also compared to the profile of mean activations, using the comparator that yielded the highest overall performance for these two methods.

#### Natural Scenes Dataset Region Identification

We first compared RIA across representational alignment measures in an fMRI dataset, where it quantifies how often each measure classifies a given brain region as more similar to instances of the same region in separate subjects than it is to instances of different regions from those subjects. The Natural Scenes Dataset^1^ (NSD) is a 7T fMRI dataset with neural responses to natural scene images collected from eight subjects, with 30-40 scanning sessions per subject. Each subject viewed up to three repetitions of 10,000 images, with 1,000 of these images shared across subjects. Since the region identification paradigm requires common stimuli across subjects, analysis was restricted to these 1,000 shared stimuli. These stimuli were further restricted to images that were viewed at least twice by all eight subjects (since computing crossnobis distance requires at least two trials per stimulus), yielding a final set of 766 stimuli. Draws of 10, 20, 50, 100, or 150 stimuli were used, using as many fully disjoint samples as possible (e.g., seven samples of 100, or 5 samples of 150).

As regions of interest (ROIs), we used 14 retinotopic and category-selective regions: retinotopic regions V1, V2, V3, and hV4; the face-selective regions FFA and OFA; the body-selective regions EBA and FBA; the word-selective regions OWFA, VWFA, and mfs-words; and the place-selective regions OPA, PPA, and RSC. Activations from these regions were extracted using the volumetric masks provided in the NSD, with no further voxel selection performed. The NSD provides single-trial beta values computed using three different methods, of which we used the ‘b3’ variant, which allows the shape of the hemodynamic response function (HRF) to vary across voxels, and uses ridge regression and the GLMdenoise^7^ technique to improve the estimates.

The cross-validated Mahalanobis (crossnobis) estimator of representational distance takes into account varying levels of noise between voxels. If multivariate noise normalization is used, it also takes into account correlated noise between voxels. Univariate noise normalization was applied for these analyses. The stimulus-conditional voxel noise covariance matrix was computed using beta values from the images that were unique to each subject (up to a maximum of 9000). We followed the approach of taking the differences between the single-trial beta values for each stimulus presentation and the mean beta across repetitions of that stimulus and using these differences to compute the covariance between voxels, rather than the alternative approach of using the timeseries residuals from the GLM.

For the whitened RDM comparators, the matrix **V** of covariances among dissimilarity estimates was computed using data from the held-out test subject. For the NSD, **V** was estimated using the heuristic of setting the between-stimulus distances to zero. We made this choice after observing that RIA for the NSD actually declined for increasing numbers of stimuli when using whitened RDM comparators when all distances were used for computing **V**, but was restored after making this adjustment. This could plausibly be due to the noisiness of the BOLD pattern estimates and the large number of stimuli we analyzed for the NSD dataset, which could exacerbate the numerical instability involved in inverting **V** (see “Using whitened RDM comparators in framed RSA” for more details).

RIA was compared between alignment measures using matched-pairs *t*-tests, with the specific stimulus draw serving as the matching variable (e.g., framed RSA vs. standard RSA computed on the same 10 stimuli). This was performed separately for each RDM comparator and each number of stimuli.

For the NSD, a further analysis was performed to examine the effects of varying the norm of **c**. Specifically, **c** was set to have norm .5*µ*, 1*µ*, or 2*µ*, where *µ* is the mean of the norms of the stimulus patterns, with accuracy being compared within each RDM comparator and number of stimuli as with the other analyses.

#### Things Ventral Stream Spiking Dataset Region Identification

We then compared RIA across representational alignment measures in a macaque neural recording dataset to characterize how well each measure could capture aspects of the neural population code that are specific to each brain region and replicable across individual animals, once more treating different instances of the same region as a proxy for the “ground-truth” computational model of that region. The Things Ventral Stream Spiking Dataset (TVSD) contains neural recording data from V1, V4, and IT cortex of two macaque monkeys.^20^ Since this dataset has two monkeys with three brain regions each, RIA here reflects how often each alignment measure labels a given brain region in one monkey as more similar to the same brain region in the other monkey than to the other two brain regions in the other monkey (e.g., how often V1 in Monkey 1 is more similar to V1 in Monkey 2 than it is to V4 and IT of Monkey 2). This dataset includes 20,000 images that were each shown once to each macaque, and 100 images that were shown 30 times to each macaque. Since crossnobis distance requires multiple repetitions of each stimulus, we used the set of 100 images.

Since only a small set of images was available for this dataset (100 images for the TVSD, versus 766 for the NSD and thousands for the DNN analysis), for our statistical tests we used the following bootstrap resampling procedure. We conducted 100 “outer” sampling iterations in which we randomly drew, with replacement, 30 trial indices from the set of 30 trials, stipulating that draws from the first 15 trials were included in the first crossvalidation fold in computing the crossnobis distance, and draws from the second 15 trials were included in the second crossvalidation fold (no stratification was applied, such that the two folds could have varying numbers of trials). Within each of these “outer” samples of trial indices, we then drew 200 “inner-loop” samples of 3, 5, 10, 20, or 50 images, allowing for image repeats across samples but not within samples. RIA was computed for each of these inner-loop samples (using the trial indices drawn in the outer loop for all stimuli) and averaged within each outer-loop iteration, yielding 100 bootstrapped RIA values for each combination of number of stimuli, RDM comparator, and alignment measure. To allow for matched-pairs t-tests, the same outer-loop trial samples were used across all RDM comparators and alignment measures (i.e., differences were always taken between RIA values computed from the same trials). The 200 inner-loop image samples were not matched across conditions in this way; since unmatched inner-loop draws add variance to the paired differences, this renders the tests conservative rather than anticonservative. Since the outer loop of this procedure samples across trials, this procedure statistically generalizes across hypothetical further trials within these recording sessions. However, it does not generalize across animals (since only two monkeys were used), neurons (since all recorded neurons from each region were always used), or stimuli (since these were only subsampled in the inner loop, and were not a source of outer-loop variability). Matched-pairs *t*-tests were then used to compare RIA between conditions, where the standard error was the standard deviation of the differences across the 100 bootstrap samples.

The channel noise covariance matrix (for whitening the pattern estimates) was computed using the differences between the beta value for each trial and the mean beta across repetitions of the image for that trial, using all 30 trials from the set of 100 repeated images. As with the NSD, univariate noise normalization was applied.

An important difference between the TVSD and the other two datasets is that there are only two “subjects” (versus eight for the NSD, and ten for the DNN dataset), such that there is no meaningful distinction between “training” and “test” subjects for each trial. To account for this, two adjustments were made to the analysis pipeline. First, accuracy was tallied in both possible “directions” on each trial. That is, whereas for the other two datasets accuracy was tallied as the proportion of trials where the held-out test RDM was more similar to the correct training RDM than to the other training RDMs (i.e., with the training and test subjects being treated asymmetrically), for the TVSD both directions were tallied. Thus, for each stimulus draw, a maximum of six “points” were possible: one point if Monkey 1’s V1 was more similar to Monkey 2’s V1 than to Monkey 2’s V4 and IT, another point if Monkey 2’s V1 was more similar to Monkey 1’s V1 than to Monkey 1’s V4 and IT—the opposite direction—and so on for the other possible comparisons. Second, for the whitened RDM comparators, instead of using the data from the “test” subject to compute **V** (the matrix of covariances among RDM entries) as with the other two datasets, **V** was computed for both monkeys, the RDM similarity was separately computed using the two estimates of **V**, and the two resulting similarity scores were averaged to derive a final score. Unlike for the NSD, in estimating **V** we used the estimated stimulus-to-stimulus distances rather than treating these distances as zero, since these distances were less noisy than for the fMRI data, and since fewer stimuli were analyzed than for the NSD, such that inverting **V** is more numerically stable.

#### Deep Neural Network Layer Identification

In the third dataset, we extend the region identification paradigm to deep neural network (DNN) layers. The analysis follows the same logic, replacing brain regions with DNN layers, and replacing individual subjects with DNN instances of the same architecture trained from different random seeds with the same task. Thus, in this case we quantify how often each representational alignment measure identifies a given DNN layer (e.g., Conv3) as being more similar to the same layer in different random seeds than it is to different layers (e.g., FC2) in different seeds. Beyond being of interest in their own right, DNNs offer several further benefits. First, they are “glass-box” systems whose ground-truth activations can be known with full precision. Second, while the DNNs used here are deterministic, we add varying levels of measurement noise in simulation so as to characterize the performance of framed RSA and the alternative methods as a function of the noise level.

Ten instances of AlexNet trained on ImageNet from different random seeds were used, with activations extracted from all layers from the second pooling layer onward (in order: Pool2, Conv3, ReLU3, Conv4, ReLU4, Conv5, ReLU5, Pool3, FC1, ReLU6, FC2, ReLU7, and FC3), for 13 layers in total. Ten disjoint sets of images were randomly drawn from MSCOCO, with a sample size of 3, 5, 10, 20, or 50 stimuli. Noise was added to the activations from each layer using the following procedure. In order to roughly equate the relative noise magnitude across layers, for each stimulus sample the grand standard deviation across stimuli and channels was computed, and isotropic noise with standard deviation set to either 1× or 10× the activation standard deviation was added to all units. Two “trials” per image were drawn in this manner. The matrix of covariances among RDM entries **V** was computed using the distances between these noisy patterns, with the **V** from the held-out test seed being used to compare RDMs. For this analysis we used Equation 34 as given to compute **V**, using all estimated distances (i.e., we did not pragmatically set the between-stimulus distances to zero when estimating **V**, as we did with the NSD). Matched-pairs *t*-tests (with the specific stimulus draw as the matching variable) were used to compare layer identification accuracy between representational alignment metrics within each RDM comparator and each number of stimuli.

For framed RSA, **c** was set to have norm equal to the mean norm of the stimulus patterns for each sample, with this norm being computed after noise was added (for parity with the other two datasets, where no “ground-truth” noiseless patterns were available).

